# An early-diverging Caladeniinae orchid reveals diversification of terpene and apocarotenoid pathways underlying floral scent evolution

**DOI:** 10.64898/2026.09.17.752494

**Authors:** Ya-Nan Zhao, James Perkins, Zemin Wang, Eran Pichersky, Rod Peakall, Fei Zhou, Darren C. J. Wong

**Affiliations:** State Key Laboratory of Pharmaceutical Biotechnology, School of Life Sciences, Nanjing University, Nanjing 210023, China; Ecology and Evolution, Research School of Biology, The Australian National University, Canberra, ACT, 2600 Australia; College of Life Science and Technology, Gansu Agricultural University, Lanzhou, China; Department of Molecular, Cellular and Developmental Biology, University of Michigan, Ann Arbor, MI 48104, USA; School of Agriculture, Food, and Wine, Adelaide University, Glen Osmond, SA 5064, Australia; Waite Research Institute, Adelaide University, Glen Osmond SA 5064

**Keywords:** Floral scent, Pollination, Terpene synthase, Carotenoid cleavage dioxygenase, Orchidaceae

## Abstract

Orchids exhibit extraordinary floral diversity, yet the molecular mechanisms underlying floral scent evolution remain poorly understood. Guided by recent phylogenomic reconstruction of the Australian orchid subtribe Caladeniinae, we used *Glossodia major*, an earlier-diverging food-deceptive lineage, as an evolutionary anchor for investigating floral volatile diversification. Scent-producing petals showed coordinated expression of terpene synthase (TPS), carotenoid and carotenoid cleavage dioxygenase (CCD) pathways. Functional characterisation identified GmTPS-b3, GmTPS-b4 and GmTPS-a1 as contributors to floral monoterpene and sesquiterpene production, while GmCCD1 produced geranylacetone, the dominant floral volatile, from ζ-carotene. GmCCD4 and GmCCD7a showed additional, partially overlapping apocarotenoid-forming activities. Integration of tissue-specific expression, enzyme activity and subcellular localisation linked these pathways to the emitted floral bouquet. Functional comparisons with characterised *Caladenia* homologues revealed biochemical conservation and catalytic and compartmental specialisation, while expression profiling across 31 species identified pollination strategy-associated regulatory divergence in *TPSa1* and *TPSb3*, consistent with corresponding shifts in terpenoid-rich floral scent. We propose that differential modification and deployment of conserved biosynthetic pathways contributed to the attenuation and re-emergence of terpenoid-rich floral scent during Caladeniinae diversification. More broadly, our findings provide candidate mechanistic links between molecular evolution, pollinator-mediated selection and floral diversification.

## Introduction

Most flowering plants rely on animal pollinators for reproductive success and have evolved diverse floral traits that signal to and attract potential flower-visiting visitors, including bees, birds, butterflies, and flies ^1,2^. These floral traits are increasingly viewed as integrated signalling systems, with combinations of colour, scent, and reward jointly shaping pollinator interactions rather than acting independently ^3^. Floral scent, in particular, represents one of the most versatile and information rich signals, mediating plant–pollinator interactions over distances exceeding those of colour alone ^4^. These volatile organic compounds (VOCs) often conveying complex information about reward availability, species identity, and phenological state, and are often integral to both pollinator attraction and foraging decisions ^5^. The diversity and complexity of the floral volatiles produced by orchids is particularly striking. Indeed, many of the more than 1,700 floral volatile compounds reported from plants occur in one or more orchid species ^6^. This chemical diversity mirroring the extraordinary diversity of floral forms, and the prevalence of specialised and deception pollination strategies within this megadiverse family ^7,8^.

Generalised food deception, in which rewardless flowers mimic the signals of rewarding species, is among the most common strategies, often involving combinations of visual and olfactory traits that exploit the foraging behaviour of generalist pollinators ^9,10^. Numerous classes of floral volatiles, including terpenoids, benzenoids, and fatty acid derivatives, have been implicated in orchid pollinator attraction ^11^, reflecting both the diverse ecological pressures imposed by different pollination strategies and the evolutionary diversification of their underlying biosynthetic pathways ^12,13^. However, despite predicted biosynthetic origins in several systems ^14–16^, the genetic and biochemical basis underlying floral volatile production has been confirmed in relatively few cases ^17–22^. Consequently, a key unresolved question is how conserved floral volatile pathways have been repeatedly modified through changes in gene expression, enzyme function, and subcellular localisation to generate the remarkable diversity of orchid floral scents.

The Australian orchid subtribe *Caladeniinae* (Orchidoideae: Diurideae) provides a powerful study system for investigating the evolution and molecular basis of floral volatile diversity. *Caladenia* sensu lato ^23–25^, is one of the most diverse Australian orchid lineages (>350 species), encompassing sexually deceptive, food deceptive, and food rewarding taxa that exhibit striking variation in flower morphology, colour and scent ^26^. For example, sexual deception in *C. plicata* is mediated by a blend of the biosynthetically distinct compounds (S)-β-citronellol and 2-hydroxy-6-methylacetophenone ^18^, whereas *C. crebra* uses unusual sulfur-containing (methylthio)phenols ^27^. By contrast, the putatively food-deceptive *C. denticulata* emits an α-pinene-dominated bouquet containing diverse lower-abundance monoterpenes and sesquiterpenes ^20^. The widespread terrestrial orchid, *Glossodia major* (syn. *Caladenia major*), occurs across a broad range of coastal and inland habitats in southeastern Australia ^28^, as an earlier-diverging sister lineage to *Caladenia*, offers a phylogenetically informative comparison for investigating the evolution of these contrasting scent chemistries. This bee pollinated species produces visually striking flowers, characterised by a white central region surrounded by mauve to bluish purple petals, with pigmentation derived from delphinidin based anthocyanins ^29,30^. In addition to its conspicuous visual display, the flowers emit a strong, sweet scent, frequently described as confectionery like, with fruity nuances. Recent work has shown that pale and white floral morphs arise from disruption of anthocyanin biosynthesis through mutations in the structural gene dihydroflavonol 4 reductase ^30^. However, the chemical and genetic basis of the strong floral scent of *G. major* remains unresolved.

As a first step towards a systems-level understanding of floral scent in *G. major*, we integrated multi-tissue transcriptomics, floral headspace profiling and functional characterisation to address four questions: (i) Which floral volatiles are emitted? (ii) Which biosynthetic pathways are enriched in scent-producing floral tissues relative to non-active vegetative tissues? (iii) To what extent does the emitted bouquet reflect tissue-specific pathways inferred from the transcriptome? and (iv) Which genes and enzymes underpin the production of these volatiles? We further asked how the expression of selected TPS orthologues varies among representative Caladeniinae lineages spanning food reward, food deception, sexual deception and inferred reversals to food deception within predominantly sexually deceptive radiations.

Through functional characterisation, we identify candidate TPS- and CCD-mediated pathways underlying two major volatile classes, terpenoids and volatile apocarotenoids, linking tissue-specific gene expression and enzyme activity to the emitted floral bouquet. Comparisons with previously characterised *Caladenia* homologues, together with broader TPS expression profiling, provide an evolutionary framework for examining floral scent transitions from earlier food-deceptive lineages through sexual deception and subsequent reversals to food deception.

## Results

### Floral volatile profile of *G. major*

Floral headspace collections of *Glossodia major* were obtained, and putative terpenoid VOCs were tentatively identified by comparing the mass spectra obtained from GC-MS against several reference spectra databases (e.g. NIST-17) and authentic standards when possible (**Fig. 1a**). The carotenoid breakdown product, geranylacetone (6,10-dimethylundeca-5,9-dien-2-one) was the dominant floral VOC, representing 30.1 ± 1.1% of the total headspace. A series of monoterpenes were also present, including α-pinene (17.7 ± 1.4%), β-pinene (4.8 ± 0.3%), 1,8-cineole (10.2 ± 0.7%), and α-terpineol (10.8 ± 0.7%). Additional monoterpenes and their oxygenated derivatives, such as limonene, β-myrcene, β-ocimene, and 6-oxo-cineole, were detected at relatively low abundances (∼0.5–2%). The sesquiterpene β-caryophyllene was the principal sesquiterpene identified, though also at low levels (∼1%). Several other compounds were present at trace quantities (< 0.5%), including other monoterpenes (e.g., sabinene, limonene oxide), sesquiterpenes (e.g., α-humulene, α-farnesene), and carotenoid breakdown products (e.g., sulcatone, dihydroactinidiolide, *(E)*-geranylacetone, β-ionone). Collectively, isoprenoid-derived volatiles, comprising monoterpenes, sesquiterpenes, and apocarotenoids, accounted for ∼81% of the total floral headspace, with phenylpropanoids/benzenoids representing a further ∼13% (**Fig. 1b**). This pronounced dominance of terpene- and apocarotenoid-derived volatiles motivated our subsequent targeted investigation into the expression and functional role of terpene synthase (TPS) and carotenoid cleavage dioxygenase (CCD) gene families in *G. major*.

**Figure 1.**
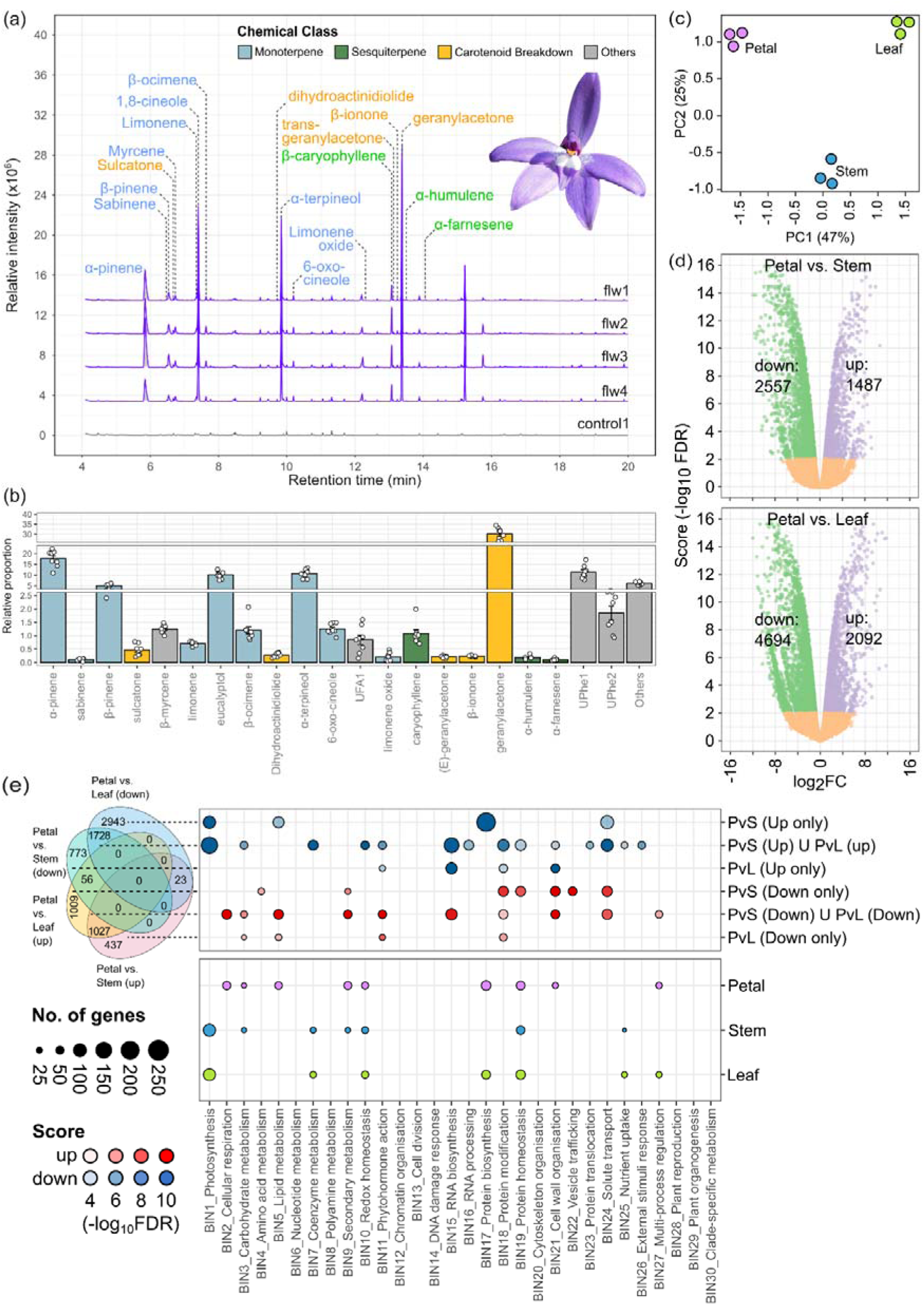
Floral volatile profile and comparative tissue transcriptomics of *Glossodia major.* (a) Representative total ion chromatograms (TICs) of floral headspace collections from *G. major* analyzed by GC–MS. Detected compounds are color-coded according to chemical class: monoterpenes (light blue), sesquiterpenes (green), apocarotenoids (dark yellow), and other compounds (grey). Chromatograms are shown for four pooled floral samples alongside an environmental control. (b) Relative abundance and estimated emission rates pg flower^-1^ h^-1^ of identified VOCs, calculated from integrated TIC peak areas after subtracting control background signals. (c) Principal component analysis (PCA) of transcriptomic profiles from petals (pink), stems (blue), and leaves (green) based on expression across 47,779 genes. (d) Volcano plots illustrating differential gene expression between scent-producing petals and non-active vegetative tissues (FDR < 0.01; upregulated in purple, downregulated in green, non-significant in orange). (e) Functional enrichment analysis of MapMan BIN v4 categories (FDR < 0.01) based on the top 1% most highly expressed genes per tissue, and unique and shared differentially expressed genes (DEGs) from Petal vs. Stem and Petal vs. Leaf contrasts. Venn diagrams summarize DEG overlaps alongside enriched functional categories (circle size and opacity denote annotated gene count and enrichment significance, respectively).

### Tissue-specific floral transcriptome comparisons and their enriched pathways

Principal component analysis of the *G. major* reference floral transcriptome (**Supplementary data S1**) showed a clear separation of samples based on tissue type (PC1: 47% and PC2: 25%), with the respective *Petal*, *Leaf* and *Stem* tissues forming distinct groups along the first two axes (**Fig. 1c**). Two complementary tissue-specific comparisons/contrasts were of special interest in this study. They were comparison of the active (floral scent-producing) petal versus non-active leaf (Petal–Leaf) and non-active stem (Petal–Stem). Of the total 47779 expressed genes, 6786 (2092 upregulated, 4694 downregulated) in the Petal–Leaf contrast and 4044 (1487 upregulated, 2557 downregulated) in the Petal–Stem contrasts were differentially expressed (DE) (FDR < 0.01) (**Fig. 1d**).

Closer inspection of the overlapping DEGs revealed that **1,027** genes were co-ordinately upregulated and **1,728** were co-ordinately downregulated across both the Petal–Leaf and Petal–Stem contrasts (**Fig. 1e**). In addition, tissue-specific differential expression was observed, whereby **437** and **1,009** genes were uniquely upregulated in Petals relative to Stem and Leaf, respectively, while **773** and **2,943** genes were uniquely downregulated in these same respective contrasts. Differentially expressed genes co-ordinately upregulated in both Petal–Leaf and Petal–Stem contrasts were enriched for several key biological processes (**Supplementary data S2**). These included pathways that were also enriched among highly expressed petal genes, such as cellular respiration (BIN2), lipid metabolism (BIN5), secondary metabolism (BIN9), and cell wall organisation (BIN21), as well as additional terms uniquely associated with DEG sets, including phytohormone action (BIN11) and RNA biosynthesis (BIN15). Conversely, enriched terms in co-ordinately downregulated genes largely aligned with enriched pathways that were highly expressed in Stem and Leaf tissues.

Furthermore, classes of genes deemed to very highly expressed (top 1%) in the vegetative Leaf and Stem tissues included those associated with photosynthesis (BIN1), coenzyme metabolism (BIN7), redox homeostasis (BIN10), protein homeostasis (BIN19), and nutrient uptake (BIN25) (**Fig. 1e, Supplementary data S3**). Conversely, the expression of genes pertaining to cellular respiration (BIN2), carbohydrate metabolism (BIN3), lipid metabolism (BIN5), secondary metabolism (BIN9), and cell wall organisation (BIN21) were enriched in petals.

### Comparative expression of candidate isoprenoid precursor biosynthesis pathways across floral and vegetative tissues

In plants, the plastidial 2-C-methyl-D-erythritol 4-phosphate (MEP) pathway and the cytosolic/endoplasmic reticulum/peroxisomal mevalonate (MVA) pathway constitute two well-established routes supplying the universal C5 isoprenoid precursors, isopentenyl diphosphate (IPP) and its isomer dimethylallyl diphosphate (DMAPP) ^31,32^. Consistent with this, secondary metabolism (BIN9) was the most significantly enriched category among concordantly upregulated genes in the Petal–Leaf and Petal–Stem contrasts. Within this category, genes associated with specialized terpenoid metabolism (BIN9.1), including the MVA pathway (BIN9.1.1), terpene biosynthesis (BIN9.1.4), and carotenoid biosynthesis (BIN9.1.6), contributed strongly to this enrichment (**Supplementary data S2**).

Candidate genes associated with the cytosolic MVA pathway included two acetyl-CoA C-acetyltransferases (*GmAACT1/2*), one 3-hydroxy-3-methylglutaryl-CoA synthase (*GmHMGS*), two 3-hydroxy-3-methylglutaryl-CoA reductases (*GmHMGR1/2*), two mevalonate kinases (*GmMVK1/2*), one phospho-mevalonate kinase (*GmPMK*), and one diphospho-mevalonate decarboxylase (*GmMPDC*). All but two genes (*GmAACT2* and *GmMVK1*) were differentially expressed (**Fig. 2, Supplementary data S4**). Notably, *GmAACT1*, *GmHMGS1*, *GmHMGR1/2*, *GmPMK*, and *GmMPDC* were coordinately upregulated in both Petal–Leaf and Petal–Stem contrasts, consistent with a potentially enhanced precursor supply through the MVA pathway. Furthermore, *GmAACT1* was among the most highly expressed transcripts in floral petals.

**Figure 2.**
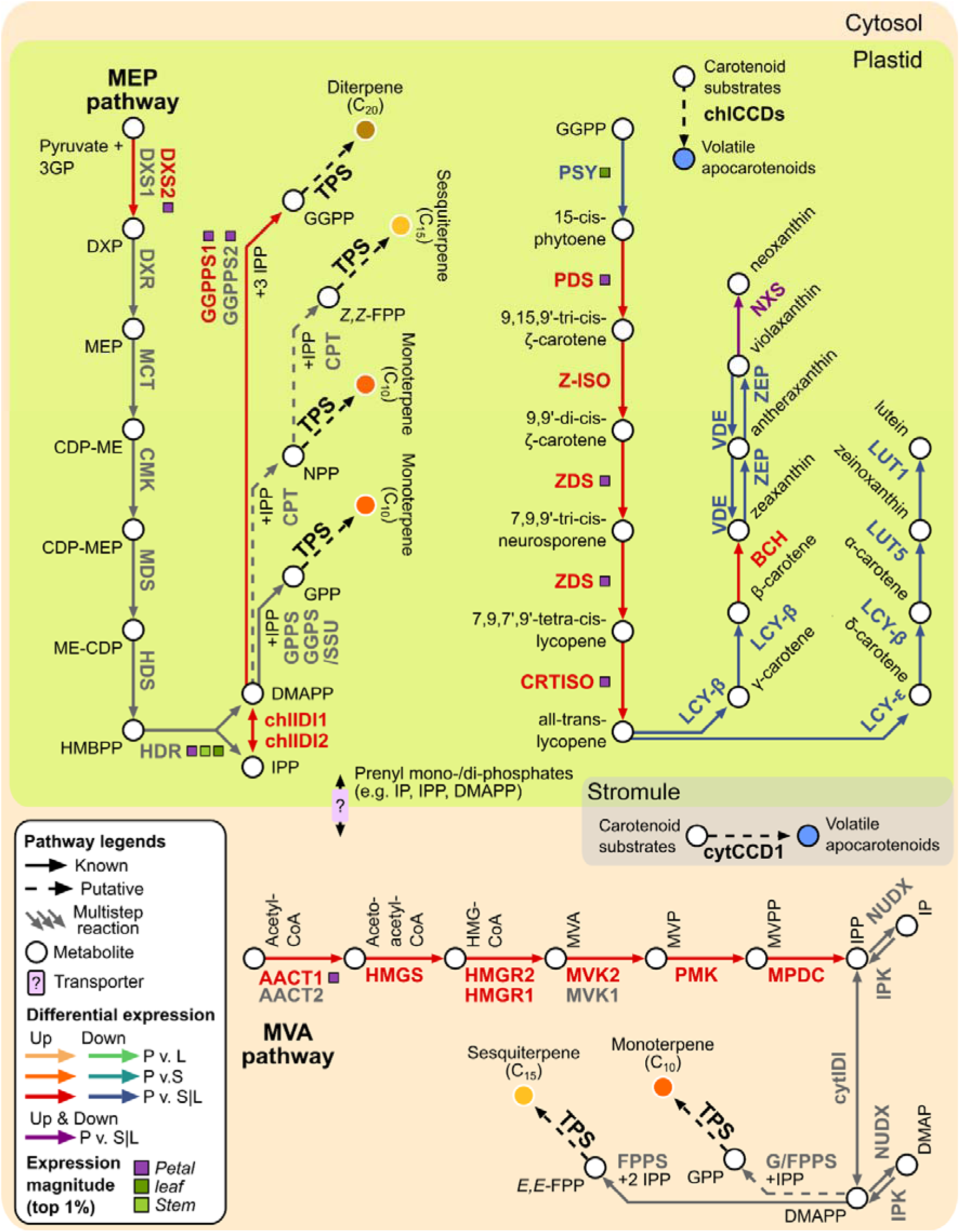
Putative terpenoid and apocarotenoid biosynthetic pathways associated with floral volatile production in *Glossodia major*. Enzymes (bold text), metabolites (circles), and reactions involved in the plastid-localized methylerythritol phosphate (MEP; green background) and cytosolic mevalonate (MVA; beige background) pathways are shown. These pathways generate the universal C_5_ isoprenoid precursors isopentenyl diphosphate (IPP) and dimethylallyl diphosphate (DMAPP), which support the biosynthesis of diverse terpenoid classes, including monoterpenes, sesquiterpenes, diterpenes, and carotenoids. plastid-localized carotenoid biosynthetic pathway from geranylgeranyl diphosphate (GGPP) to The the xanthophylls lutein and neoxanthin is also depicted, highlighting potential carotenoid substrates for volatile apocarotenoid production. Putative carotenoid cleavage reactions mediated by carotenoid cleavage dioxygenases (CCDs) are indicated within plastids, whereas cytosolic CCD1-mediated cleavage reactions are shown separately. A stromule extension highlights a putative connection between plastid and cytosolic carotenoid metabolism, through which carotenoid cleavage products may become accessible to cytosolic CCD1. Coloured arrows indicate significant differential expression between scent-producing petals and vegetative tissues (Petal vs Stem and/or Petal vs Leaf), with shades of orange-to-red and green-to-blue denoting upregulation and downregulation, respectively (FDR < 0.01), purple indicating inconsistent regulation between comparisons, and grey indicating no significant differential expression. Coloured squares denote genes whose transcript abundance ranked within the top 1% of expressed genes in the corresponding tissue transcriptome. **Abbreviations: MVA pathway**: AACT, acetyl-CoA C-acetyltransferase; HMGS, 3-hydroxy-3-methylglutaryl-CoA synthase; HMGR, 3-hydroxy-3-methylglutaryl-CoA reductase; MVK, mevalonate kinase; PMK, phosphomevalonate kinase; MPDC, mevalonate diphosphate decarboxylase. **MEP pathway**: DXS, 1-deoxy-D-xylulose 5-phosphate synthase; DXR, 1-deoxy-D-xylulose 5-phosphate reductoisomerase; MCT, 2-C-methyl-D-erythritol 4-phosphate cytidylyltransferase; CMK, 4-(cytidine 5′-diphospho)-2-C-methyl-D-erythritol kinase; MDS, 2-C-methyl-D-erythritol 2,4-cyclodiphosphate synthase; HDS, 4-hydroxy-3-methylbut-2-enyl diphosphate synthase; HDR, 4-hydroxy-3-methylbut-2-enyl diphosphate reductase. IPP and DMAPP metabolism: IPK, isopentenyl phosphate kinase; NUDX, NUDIX hydrolase; cytIDI, cytosolic isopentenyl diphosphate isomerase; chlIDI, chloroplastic isopentenyl diphosphate isomerase; G/FPPS, geranyl/farnesyl diphosphate synthase; GPPS, geranyl diphosphate synthase; GGPPS, geranylgeranyl diphosphate synthase; SSU I, type I small subunit of GPPS; SSU II, type II small subunit of GGPPS; CPT, cis-prenyltransferase. **Carotenoid pathway:** PSY, phytoene synthase; PDS, phytoene desaturase; Z-ISO, ζ-carotene isomerase; ZDS, ζ-carotene desaturase; CRTISO, carotenoid isomerase; LCYB, lycopene β-cyclase; LCYE, lycopene ε-cyclase; BCH, β-carotene hydroxylase; CYP97A, carotenoid ε-ring hydroxylase; CYP97C, carotenoid β-ring hydroxylase of the α-carotene branch; ZEP, zeaxanthin epoxidase; NSY, neoxanthin synthase; CCD, carotenoid cleavage dioxygenase.

Candidate genes associated with the plastidial MEP pathway, which supplies isoprenoid precursors for plastid-localised terpenoid biosynthesis, included two 1-deoxy-D-xylulose 5-phosphate synthases (*GmDXS1/2*), one DXP reductoisomerase (*GmDXR*), one 2-C-methyl-D-erythritol 4-phosphate cytidylyltransferase (*GmMCT*), one 4-(cytidine 5′-diphospho)-2-C-methyl-D-erythritol kinase (*GmCMK*), one 2-C-methyl-D-erythritol 2,4-cyclodiphosphate synthase (*GmMDS*), one 4-hydroxy-3-methylbut-2-enyl diphosphate synthase (*GmHDS*), and one HMBPP reductase (*GmHDR*) (**Fig. 2; Supplementary data S4**). Of these, only *GmDXS2* was coordinately upregulated in both Petal–Leaf and Petal–Stem contrasts. Notably, *GmHDR* was among the most highly expressed genes across leaf, stem, and petal tissues, while *GmDXS2* also showed elevated expression in petals.

IPP and DMAPP, generated by the MVA and MEP pathways, are interconvertible metabolites, with their equilibrium primarily maintained by IPP isomerase (IDI), which localises to multiple subcellular compartments including plastids, mitochondria, and peroxisomes. Recent studies have also implicated IP kinase (IPK) and NUDIX hydrolases (NUDX) in modulating the cellular pools of these intermediates through phosphorylation and dephosphorylation reactions, respectively ^32^. Consistent with this complexity, we identified two predicted chloroplast localised IDIs (GmchlIDI1/2), one cytosolic IDI (GmcytIDI), one IPK (GmIPK), and one cytosolic NUDX (GmNUDX1). Notably, *GmchlIDI1/2* were co-ordinately upregulated, whereas *GmNUDX1* was downregulated, in both Petal–Leaf and Petal–Stem contrasts (**Fig. 2**).

In plants, the synthesis of short-chain prenyl diphosphates is compartmentalised, with geranyl diphosphate (GPP) primarily produced in plastids, farnesyl diphosphate (FPP) in the cytosol, and geranylgeranyl diphosphate (GGPP) across multiple subcellular compartments. These intermediates are generated by short-chain prenyl diphosphate synthases (e.g., GPPS, FPPS, GGPPS), which catalyse the sequential addition of IPP units to DMAPP. Consistent with this, we identified candidate genes encoding the Arabidopsis homologues of the large (GmGGPPS1/2_LSU) and small (GmGGPPS_SSU) subunits of the heterodimeric GGPP synthase in *G. major*. In addition, a predicted cytosolic bifunctional GPP/FPP synthase homologue (GmG/FPPS) was identified. Of these, only *GmGGPPS1_LSU* was coordinately upregulated in both Petal–Leaf and Petal–Stem contrasts (**Fig. 2**). Notably, both *GmGGPPS1/2_LSU* were among the most highly expressed transcripts in petal tissues.

### Comparative expression of candidate terpene synthase genes across floral and vegetative tissues

Terpene synthases (TPS) in plants are broadly classified into at least eight subfamilies (TPS-a to TPS-h) based on phylogenetic relationships, sequence similarity, and functional diversification ^33,34^. Phylogenetic placement of candidate *G. major* TPS alongside functionally characterised TPS from a diverse set of angiosperms—including closely related *Caladenia denticulata* TPS—enabled assignment of these genes to established TPS clades. Specifically, GmTPS-c1 clustered within the TPS-c clade, GmTPS-e/f1 within the TPS-e/f clade, GmTPS-a1 within the TPS-a clade, and GmTPS-b3 and GmTPS-b4 within the TPS-b clade (**Fig. 3a, Supplementary data S6**).

**Figure 3.**
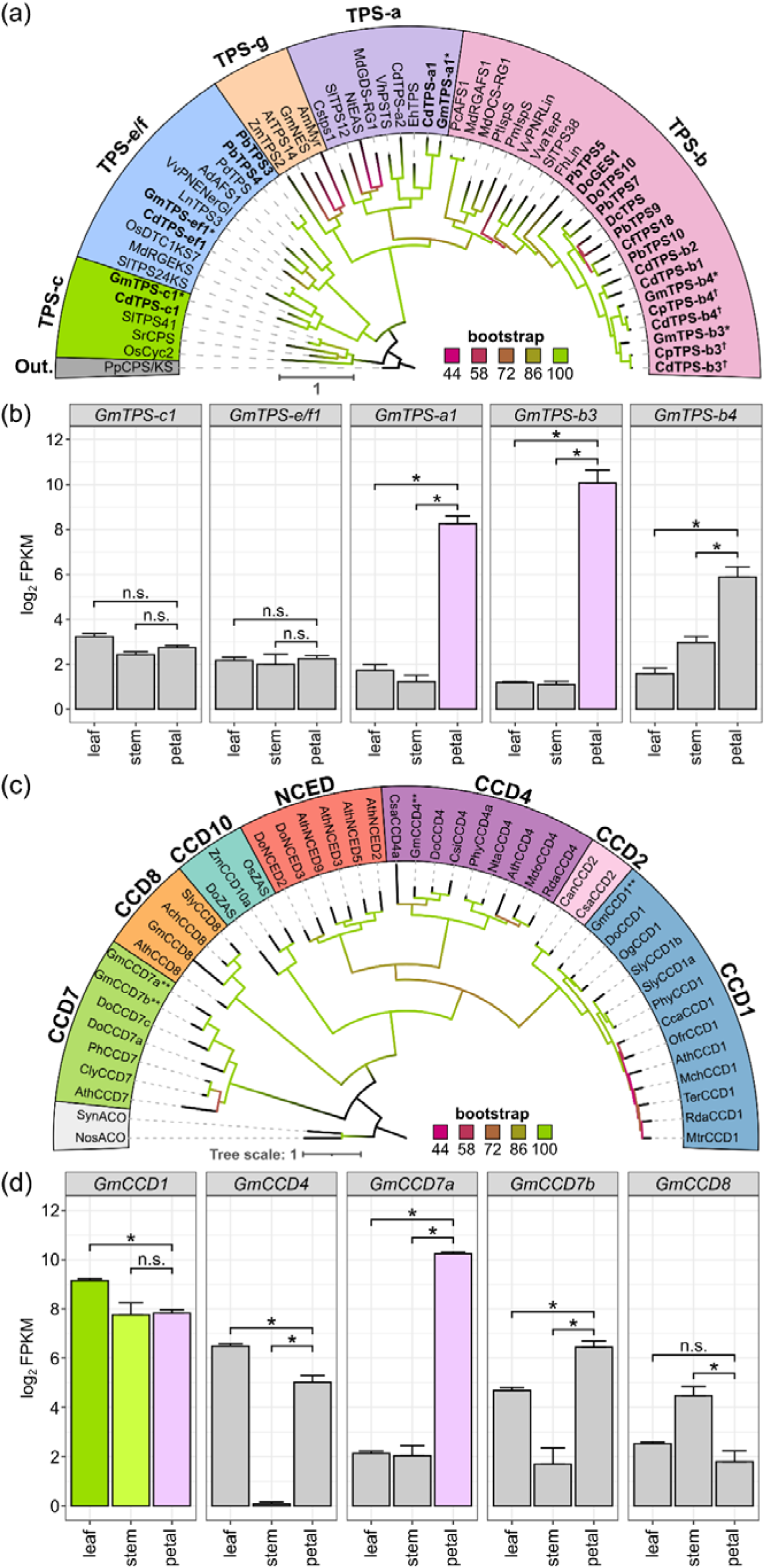
Terpene synthase (TPS) and carotenoid cleavage dioxygenase (CCD) gene families in Glossodia major. (a) Maximum-likelihood phylogeny of TPS proteins inferred using IQ-TREE from putative *G. major* TPSs and functionally characterized TPSs from diverse angiosperms, including orchids and representatives from the closely related species *Caladenia denticulata* and *C. plicata*. Shaded regions indicate TPS-c (green), TPS-e/f (blue), TPS-g (yellow), TPS-a (purple), and TPS-b (salmon pink) subfamilies; grey denotes the outgroup. Asterisks indicate putative *G. major* TPSs. The scale bar represents amino acid substitutions per site. See **Supplementary Data S6** for accession numbers and associated references. (b) Transcript abundance of putative *G. major TPS* genes in scent-producing petals and vegetative tissues (stems and leaves). Gene expression is shown as log_2_ FPKM values. Asterisks indicate genes significantly upregulated in petals relative to one or both vegetative tissues (Petal vs Stem and/or Petal vs Leaf; FDR < 0.01); *n.s.* indicates no significant differential expression. (c) Maximum-likelihood phylogeny of CCD proteins inferred using IQ-TREE from putative *G. major* CCDs and functionally characterized CCDs from diverse angiosperms, including orchids. Shaded regions indicate CCD1 (light blue), CCD2 (pink), CCD4 (lavender), CCD7 (light green), CCD8 (orange), CCD10 (turquoise), and NCED (soft red) subfamilies; grey denotes the outgroup. Asterisks indicate putative *G. major* CCDs. The scale bar represents amino acid substitutions per site. See **Supplementary Data S7** for accession numbers and associated references. (d) Transcript abundance of putative *G. major* CCD genes in scent-producing petals and vegetative tissues (stems and leaves). Gene expression is shown as FPKM values. Orange bars marked with asterisks indicate genes significantly upregulated in petals relative to one or both vegetative tissues (Petal vs Stem and/or Petal vs Leaf; FDR < 0.01), whereas *n.s.* indicates no significant differential expression. Coloured borders denote genes whose transcript abundance ranked within the top 1% of expressed genes in the corresponding tissue transcriptome.

Expression profiling revealed clear tissue-specific patterns among these candidates. *GmTPS-a1*, *GmTPS-b3*, and *GmTPS-b4* were consistently upregulated in petals relative to leaf and stem tissues, with *GmTPS-a1* and *GmTPS-b3* ranking among the most highly expressed transcripts in petals. In contrast, *GmTPS-c1* and *GmTPS-e/f1* exhibited uniformly low expression across all tissues examined. Taken together, these results identify GmTPS-a1, GmTPS-b3, and GmTPS-b4 as strong candidates contributing to mono- and sesquiterpene biosynthesis in *G. major* flowers (**Fig. 3b**).

### Comparative expression of carotenoid and apocarotenoid biosynthesis pathways across floral and vegetative tissues

Phytoene is synthesised from two molecules of geranylgeranyl diphosphate (GGPP) by phytoene synthase (PSY), and is subsequently desaturated in a stepwise manner to lycopene through the sequential action of phytoene desaturase (PDS) and ζ carotene desaturase (ZDS), via the intermediates phytofluene, ζ carotene, and neurosporene ^35,36^. These intermediates are maintained in the appropriate *trans* configuration by ζ carotene isomerase (*Z* ISO) and carotene isomerase (CRTISO). Candidate genes included one phytoene synthase (*GmPSY*), one 15-*cis*-phytoene desaturase (*GmPDS*), one 15-*cis*-ζ-carotene isomerase (*GmZISO*), one 9,9’-di-*cis*-ζ-carotene desaturase (*GmZDS*), and one carotenoid isomerase (*GmCRTISO*). All but *GmPSY* were co-ordinately upregulated in both Petal–Leaf and Petal– Stem contrasts. Additionally, *GmPDS*, *GmZDS*, *GmCRTISO* were among the most highly expressed genes in the Petal tissues (**Fig. 2, Supplementary data S5**).

Lycopene is cyclised to β carotene by lycopene β cyclase (LβCY) via γ carotene, or to α carotene through the combined activity of lycopene ε cyclase (LεCY) and lycopene β cyclase (LβCY) via δ carotene. These carotene backbones are subsequently hydroxylated to form xanthophylls: β carotene is converted to zeaxanthin by β carotene hydroxylase (BCH), while α carotene is sequentially hydroxylated by β carotene hydroxylase (LUT5) and ε carotene hydroxylase (LUT1) to produce lutein. Zeaxanthin undergoes reversible epoxidation to violaxanthin via zeaxanthin epoxidase (ZEP) and violaxanthin de epoxidase (VDE), and violaxanthin is further converted to neoxanthin by neoxanthin synthase (NXS) ^35,36^. Candidate genes include one lycopene β-cyclase (*GmLCY-*β), one lycopene ε-cyclase (*GmLCY-*ε), one β-carotene hydroxylase (*GmBCH*), one ε-carotene hydroxylase (*GmCHY-*ε), one Zeaxanthin epoxidase (*GmZEP*), one violaxanthin de-epoxidase (*GmVDE*), one β-ring carotene hydroxylase (*GmLUT5*), one ε-ring carotene hydroxylase (*GmLUT1*), and several neoxanthin synthase (*GmNXS*) (**Fig 2., Supplementary data S5**). Interestingly, unlike genes associated with upstream carotenoid precursor biosynthesis, *GmLCY-* β, *GmLCY-*ε, *GmZEP*, *GmVDE*, *GmLUT5*, and *GmLUT1* were co-ordinately downregulated in both the Petal–Leaf and Petal–Stem contrasts. The primary exception was *GmBCH*, which exhibited coordinated upregulation in both comparisons, whereas *GmNXS* homologues showed no consistent pattern of differential expression.

### Comparative expression of candidate carotenoid cleavage dioxygenase genes across floral and vegetative tissues

Carotenoid cleavage dioxygenases (CCDs) constitute a functionally diverse enzyme family that catalyse position specific oxidative cleavage of carotenoid substrates, generating apocarotenoids with roles in volatile production, pigmentation, and hormone biosynthesis. In plants, CCDs are broadly classified into at least seven subfamilies (CCD1, CCD2, CCD4, CCD7, CCD8, CCD10, and NCED) based on phylogenetic relationships, sequence similarity, and functional diversification ^35,36^. Phylogenetic placement of candidate *G. major* CCDs alongside functionally characterised CCDs from a diverse set of angiosperms, including orchid representatives such as *Cymbidium*, *Oncidium*, and *Dendrobium*, enabled their assignment to established CCD clades. Specifically, GmCCD1 grouped within the CCD1 clade, GmCCD4 within CCD4, GmCCD7a and GmCCD7b within CCD7, and GmCCD8 within CCD8 (**Fig. 3c; Supplementary data S7**).

Expression profiling revealed distinct, tissue-specific regulatory patterns among these candidates. *GmCCD1* was among the most highly expressed genes across all tissues, albeit with significantly higher expression in leaf compared to stem and petal. *GmCCD4* was upregulated in petals relative to stems, but showed highest expression in leaf tissue. In contrast, *GmCCD7a* and *GmCCD7b* displayed coordinated upregulation in petals relative to both leaf and stem, with *GmCCD7a* also ranking among the most highly expressed transcripts in petal tissue. Conversely, *GmCCD8* expression was enriched in stem, with comparatively low and uniform expression in leaf and petal tissues (**Fig. 3d**). Taken together, these expression patterns suggest functional partitioning within the CCD family, with GmCCD1, GmCCD4, GmCCD7a, GmCCD7b representing strong candidates underpinning apocarotenoid biosynthesis in *G. major* flowers.

### Subcellular localization of *G. major* TPS and CCD proteins

To further resolve the spatial organisation of candidate terpene and apocarotenoid biosynthetic enzymes, we investigated their subcellular localisation using transient expression assays (**Fig. 4**). Full-length coding sequences of three TPS and four CCDs proteins were fused to the N terminus of YFP and expressed in *Nicotiana benthamiana* leaves. Confocal microscopy revealed distinct localisation patterns among CCD family members. GmCCD1–YFP was localised to the cytosol, whereas GmCCD4–YFP, GmCCD7a–YFP, and GmCCD7b–YFP exhibited fluorescence restricted to plastids. These localisation patterns are consistent with functional partitioning of CCD activity between cytosolic volatile formation and plastidial carotenoid turnover. Conversely, TPS proteins exhibited greater flexibility in subcellular localisation. Both GmTPS b3–YFP and GmTPS b4–YFP displayed dual plastid and cytosolic fluorescence signals (**Fig. 4**). Notably, the cytosolic signal, particularly for GmTPS b4–YFP, frequently appeared punctate or speckled, suggesting potential association with subcellular structures or microdomains. In contrast, GmTPS a1–YFP was confined to the cytosol, consistent with a role in sesquiterpene biosynthesis.

**Figure 4.**
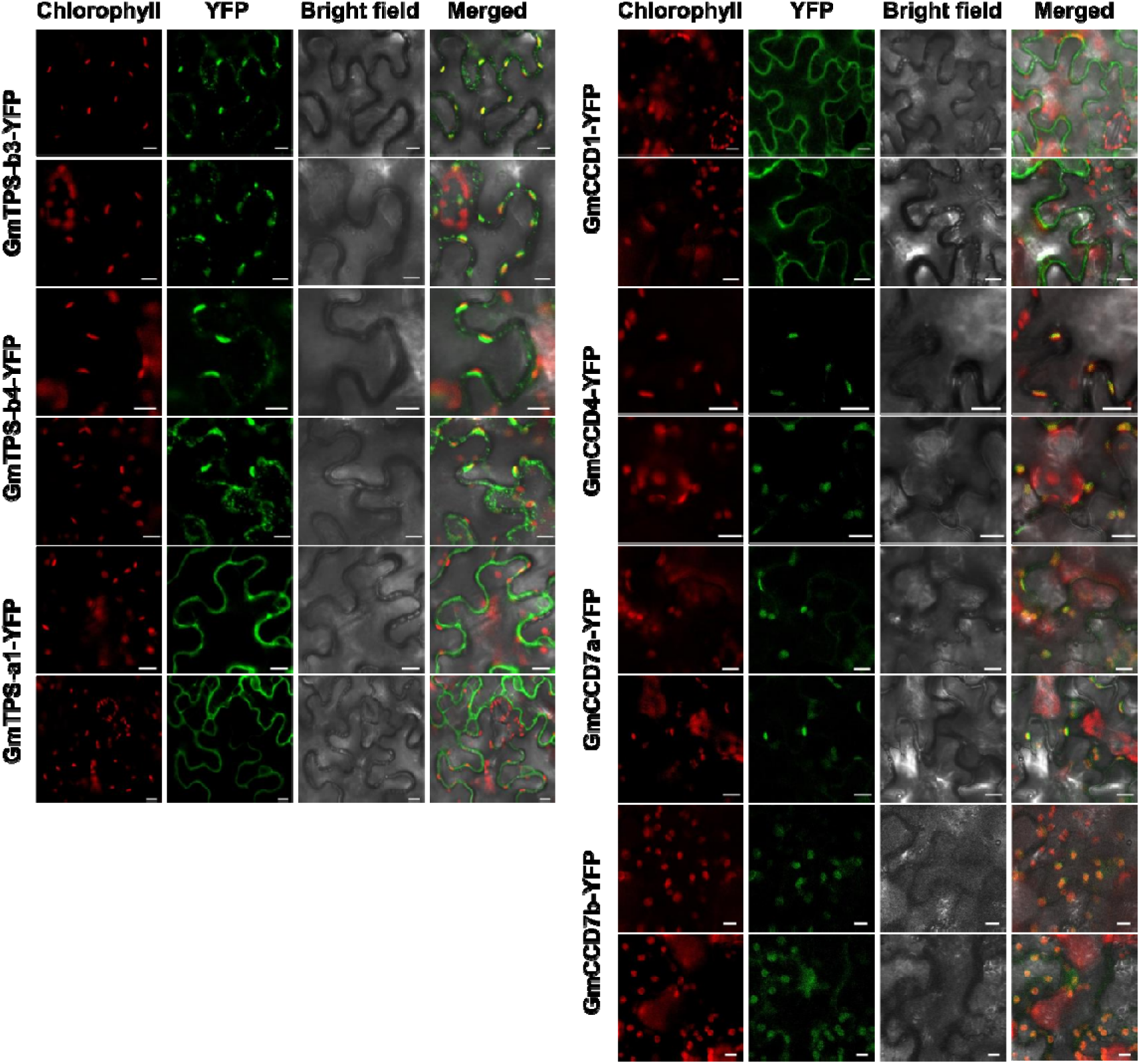
Subcellular localization of candidate terpene synthase (TPS) and carotenoid cleavage dioxygenase (CCD) proteins from *Glossodia major*. The N-terminal region (first 100 amino acids) of GmTPS-a1, GmTPS-b3, GmTPS-b4, GmCCD1, GmCCD4, GmCCD7a, and GmCCD7b was fused to YFP and transiently expressed in *Nicotiana benthamiana* leaf epidermal cells. An unfused YFP construct served as a control. Fluorescence signals were visualized by confocal laser scanning microscopy. Images show chlorophyll autofluorescence (red), YFP fluorescence (green), bright-field images (grey), and merged channels (overlay), respectively. Scale bars = 10 μm.

### Functional characterization of TPS proteins *in vitro*

To elucidate the catalytic functions of candidate TPS enzymes, purified recombinant proteins were assayed *in vitro* using a range of prenyl diphosphate substrates, including GPP, NPP, *E*,*E*-FPP, and *Z*,*Z*-FPP (**Fig. 5**). These assays revealed distinct substrate preferences and product profiles among the three GmTPS enzymes, highlighting their contributions to terpenoid diversity (**Fig. 1a**). Among the monoterpene substrates, GmTPS b3 utilised both GPP and NPP to produce a mixture of α pinene (peak 1), sabinene (peak 2), β pinene (peak 3), and β myrcene (peak 4). GmTPS b4 exhibited a more restricted product profile, converting GPP to β myrcene (peak 4), β ocimene (peak 6), and α terpineol (peak 7), while NPP supported the formation of α terpineol only (**Fig. 5a, Supplementary data S9**). In contrast, GmTPS a1 did not generate detectable monoterpene products from GPP but produced D limonene (peak 8) and (+)- 4 carene (peak 9) when supplied with NPP, indicating substrate-specific activity within the monoterpene pool.

**Figure 5.**
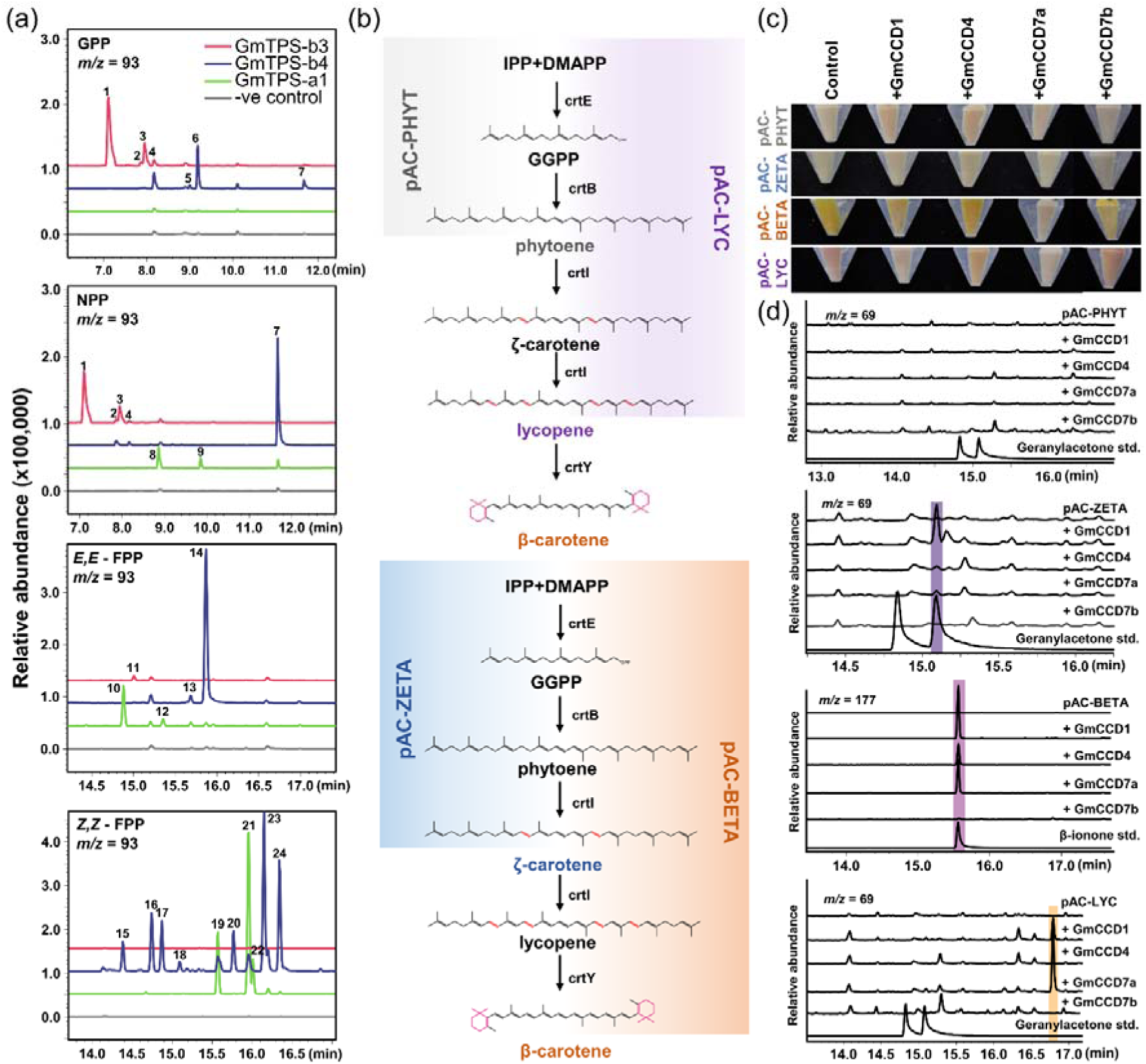
Functional characterization of *Glossodia major* terpene synthase and carotenoid cleavage dioxygenases (CCDs) enzymes. (a) GC–MS analysis of volatile products generated by recombinant *Glossodia major* TPS enzymes *in vitro*. Purified TPS proteins were incubated with GPP, NPP, *E,E*-FPP, and *Z,Z*-FPP, and reaction products were analysed by GC**–**MS as described in the Materials and Methods. The extracted ion chromatogram (EIC) at *m/z* 93 was monitored to detect terpene products. Compound identities were assigned by comparison of mass spectra and retention indices with authentic standards and/or NIST library entries. Numbered peaks correspond to: 1, α-pinene; 2, sabinene; 3, β-pinene; 4, β-myrcene; 5, unknown; 6, β-ocimene; 7, _α_-terpineol; 8, D-limonene; 9, (+)-4-carene; 10, caryophyllene; 11, cis-α-bergamotene; 12, α-humulene; 13– 14, α-farnesene isomers; 15, 7-epi-sesquithujene; 16, trans-α-bergamotene; 17, β-farnesene; 18, unknown; 19, β-curcumene; 20, α-zingiberene; 21, β-bisabolene; 22, unknown; 23, β-sesquiphellandrene; and 24, _α_-bisabolene. Mass spectra of all reaction products are provided in **Supplementary Data S9**. (b) Schematic of the carotenoid-producing *E. coli* assay system. Recombinant strains harbouring pAC-PHY, pAC-ZETA, pAC-LYC, or pAC-BETA accumulate phytoene, ζ-carotene, lycopene, and β-carotene, respectively, and were co-transformed with constructs expressing GmCCD1, GmCCD4, GmCCD7a, or GmCCD7b to assess carotenoid cleavage activity. (c) Representative phenotypes of recombinant *E. coli* cell pellets. Expression of active GmCCD enzymes resulted in visibly reduced carotenoid pigmentation relative to the corresponding carotenoid-producing controls, particularly in lycopene- and β-carotene-accumulating strains, consistent with enzymatic cleavage of intracellular carotenoid substrates. (d) GC–MS chromatograms of volatile apocarotenoid products detected from carotenoid-producing *E. coli* strains expressing individual GmCCD enzymes. Chromatograms are shown for strains accumulating phytoene, ζ-carotene, and β-carotene or lycopene. Extracted ion chromatograms (EICs) at m/z 69 and m/z 177 were monitored for the detection of volatile apocarotenoid cleavage products. Authentic β-ionone and geranylacetone standards are shown for comparison. Product identities were assigned by comparison of retention times and mass spectra with authentic standards and NIST reference spectra.

When incubated with sesquiterpene substrates, GmTPS b3 catalysed the formation of cis α bergamotene (peak 11) from *E*,*E*-FPP, although no products were detected with *Z*,*Z*-FPP. In contrast, both GmTPS b4 and GmTPS a1 generated diverse sesquiterpene profiles from both *E*,*E*-FPP and *Z*,*Z*-FPP (**Fig. 5a, Supplementary data S9**). Using *E*,*E*-FPP, GmTPS b4 produced *Z*,*E*-α farnesene (peak 13) and *E*,*E*-α farnesene (peak 14), whereas *Z*,*Z*-FPP resulted in a broader range of sesquiterpenes, including 7 *epi* sesquithujene (peak 15), *trans* α bergamotene (peak 16), β farnesene (peak 17), a β sesquiphellandrene like compound (peak 18), β curcumene (peak 19), α zingiberene (peak 20), β bisabolene (peak 21), an α bergamotene like compound (peak 22), β sesquiphellandrene (peak 23), and α bisabolene (peak 24). Similarly, GmTPS a1 exhibited robust sesquiterpene synthase activity, producing caryophyllene (peak 10) and humulene (peak 12) from *E*,*E*-FPP, and β curcumene (peak 19), β bisabolene (peak 21), and an α bergamotene like product (peak 22) from *Z*,*Z*-FPP.

Consistent with these enzymatic activities, transient expression assays demonstrated that GmTPS a1 localises to the cytosol (**Fig. 4**), in line with its capacity to utilise FPP substrates for sesquiterpene biosynthesis. None of the three TPS enzymes produced detectable products when supplied with GGPP, indicating that these enzymes function as mono and sesquiterpene synthases rather than diterpene synthases. Taken together, these results demonstrate that GmTPS b3, GmTPS b4, and GmTPS a1 exhibit distinct substrate specificities and catalytic profiles, collectively contributing to the structural diversity of mono and sesquiterpenes in *G. major*.

### Functional characterization of Carotenoid Cleavage Dioxygenase proteins *in vitro*

To functionally characterise the catalytic activity of GmCCD enzymes, we first analysed their sequence features and predicted subcellular targeting using *in silico* tools. For plastid-localised CCDs, predicted transit peptides were removed prior to heterologous expression in *Escherichia coli* to facilitate proper protein production. CCD activity was then assessed using engineered *E. coli* strains capable of accumulating defined carotenoid intermediates ^37^, including phytoene (pAC-PHYT), ζ-carotene (pAC-ZETA), lycopene (pAC-LYC), and β-carotene (pAC-BETA) (**Fig. 5b**). A two-step transformation strategy was employed to improve transformation efficiency, as described in the Materials and Methods. Following CCD expression, changes in bacterial pellet colour provided an initial qualitative indication of carotenoid cleavage (**Fig. 5c**).

To resolve reaction products, volatile compounds were extracted and analysed by GC–MS. The extracted ion chromatogram (EIC) at *m*/*z* 69 was used to detect geranylacetone, a diagnostic product of C9–C10 (C9′–C10′) cleavage of linear carotenoids such as phytoene and ζ-carotene (Yoo et al., 2023). Among the CCDs tested, only GmCCD1 catalysed cleavage of ζ-carotene to generate geranylacetone (**Fig. 5d**), whereas no activity toward phytoene was detected for this enzyme. In contrast, β-carotene served as a substrate for multiple CCDs. GmCCD1, GmCCD4, and GmCCD7a all produced β-ionone, indicating conserved cleavage activity toward cyclic carotenoids. A compound tentatively identified as pseudoionone based on mass spectral similarity to the NIST library was detected in lycopene-accumulating strains expressing GmCCD1 and GmCCD7a. This compound is consistent with C9-C10 (C9’-C10’) cleavage of lycopene. No detectable cleavage products were observed for GmCCD7b across the tested substrates.

Collectively, these results demonstrate that GmCCD1 exhibits selective activity toward ζ-carotene and broad activity toward cyclic carotenoids, while GmCCD4 and GmCCD7a preferentially cleave β-carotene. The shared production of β-ionone and pseudoionone across multiple CCDs highlights both overlapping and substrate-specific catalytic functions, contributing to the diversity of apocarotenoid products in *G. major*.

### Pollination strategy-associated divergence in TPS expression across Caladeniinae

To examine whether the contrasting expression patterns observed among functionally characterised TPS homologues extend across Caladeniinae, we compared *TPSa1*, *TPSb3* and *TPSb4* orthologue expression across 31 species using a recent phylogenomic framework (O’Donnell et al., 2025) (**Fig. 6a**). The sampling represented food reward in the earliest-diverging *Leptoceras* lineage (FR, *n* = 1), early-diverging food-deceptive lineages (FD, *n* = 12), predominantly sexually deceptive lineages (SD, *n* = 11), and subsequent shifts to food deception within predominantly sexually deceptive radiations (R, *n* = 7). Expression differed among FD, SD and R species for *TPSa1* (*H*₂ = 10.7, *P* = 0.00486) and *TPSb3* (*H*₂ = 10.1, *P* = 0.00645), but not *TPSb4* (*H*₂ = 4.25, *P* = 0.120; Fig. 7b). *TPSa1* and *TPSb3* expression was lower in SD than FD species (*P*adj = 0.00389 and 0.00927, respectively), whereas *TPSb3* expression was higher in R than SD species (*P*adj = 0.0362) (**Fig. 6b**). The species-level expression heatmap and complete statistical results are provided in **Supplementary Data S11.**

**Figure 6.**
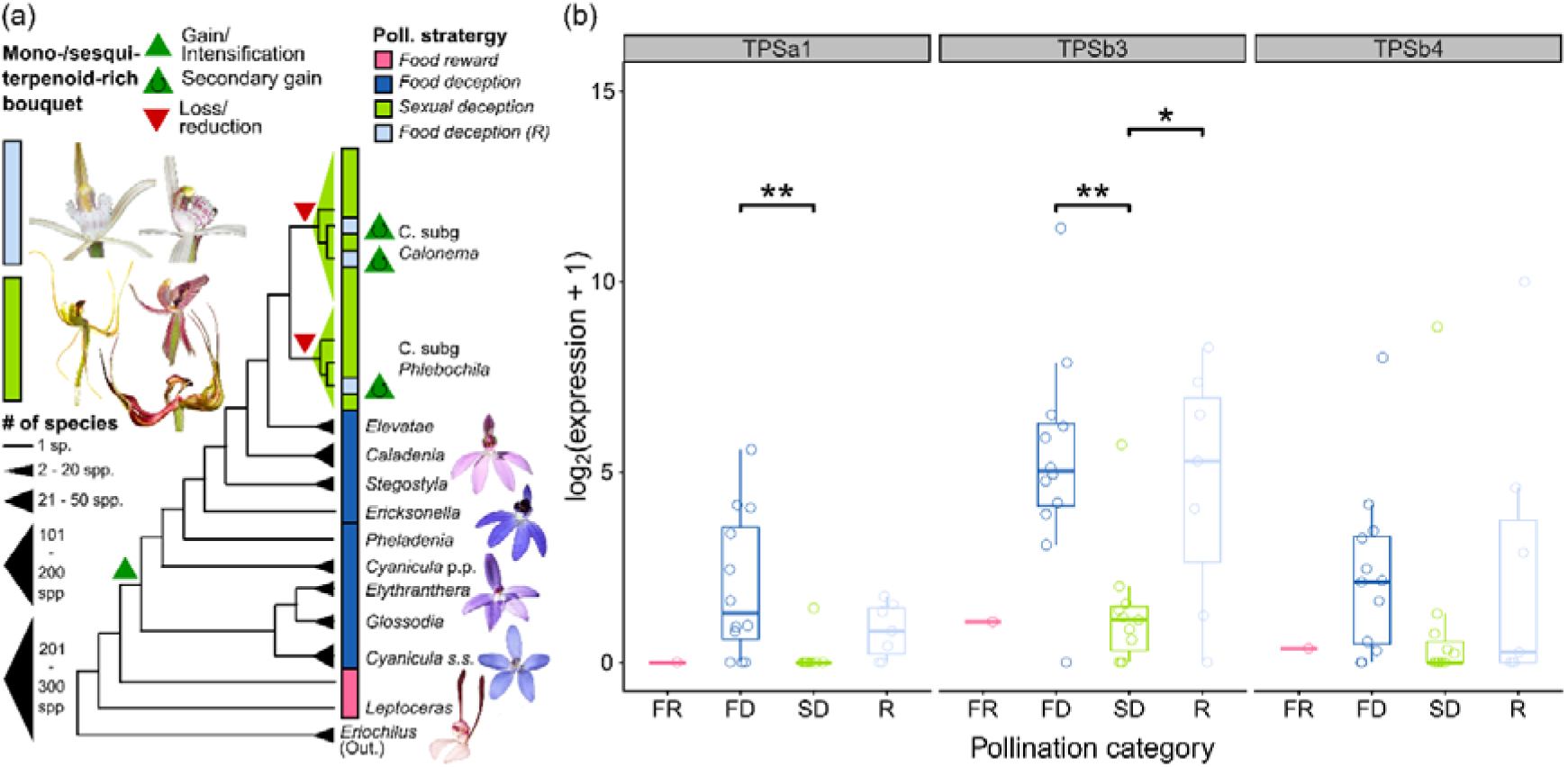
Floral terpenoid expression between pollination strategies in Caladeniinae. (a) Representative phylogeny of Caladeniinae showing major subgenera and lineages, with triangle sizes proportional to clade species richness, particularly within the species-rich subgenera *Calonema* and *Phlebochilus*. Coloured bars denote pollination strategies: food reward in the earliest-diverging *Leptoceras* lineage (FR; pink), food deception (FD; dark blue), sexual deception (SD; green), and inferred reve sals from sexual deception to food deception (R). Evolutionary changes in terpenoid-rich floral scent are indicated by symbols: a green triangle denotes intensification, a red triangle denotes severe reduction, and a green triangle with a circular arrow denotes secondary gain. The inferred reduction in monoterpenoids, sesquiterpenoids and potentially volatile apocarotenoids coincides with the radiation of predominantly weakly scented or apparently scentless SD species. The inferred re-emergence of terpenoid-rich floral scent coincides with independent shifts to food deception. (b) Interspecific expression of *TPSa1*, *TPSb3* and *TPSb4* across pollination categories. Points represent species; boxes show medians and interquartile ranges. FR was retained as the earliest-diverging reference but excluded from statistical testing because it was represented by one species. Differences were assessed using Kruskal–Wallis tests followed by Dunn’s tests with Holm correction. Only significant comparisons are shown: \**P*adj < 0.05; \*\**P*adj < 0.01. The species-level heatmap, complete statistics and underlying expression values are provided in Supplementary Data S11.

**Figure 7.**
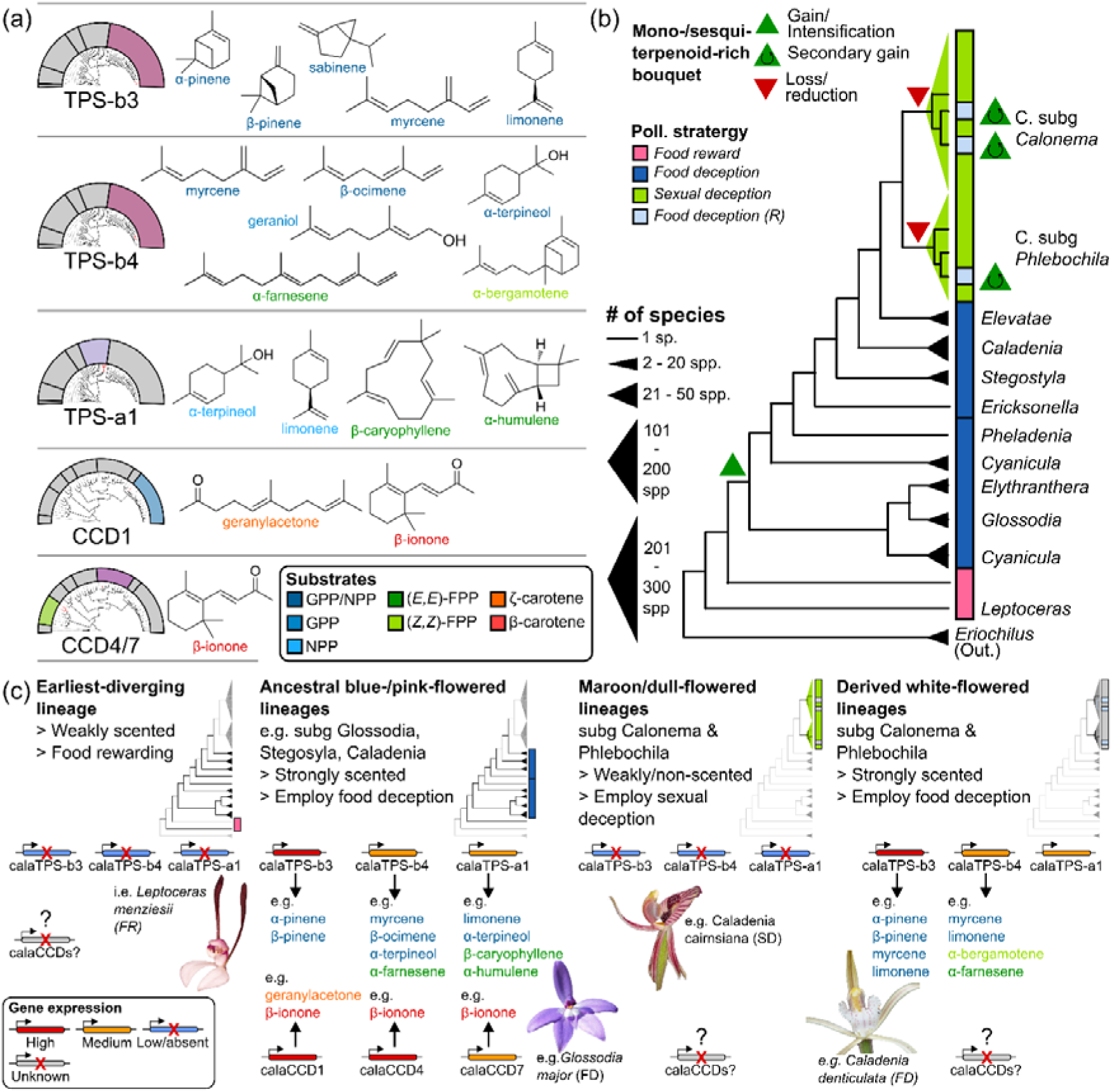
Conceptual model of terpene and apocarotenoid biosynthesis underlying floral scent diversification in the Caladeniinae. (a) Summary of functionally characterized terpene synthases (TPSs) and carotenoid cleavage dioxygenases (CCDs) from representative Caladeniinae species. Experimentally validated enzyme products detected in floral headspace samples or tissue extracts are shown alongside their corresponding enzyme classes (e.g. TPS-b3, TPS-b4, TPS-a1, CCD1, CCD4, and CCD7). Product colours correspond to their confirmed precursor substrates, including GPP/NPP-derived monoterpenes, *E,E*-FPP- and *Z,Z*-FPP- derived sesquiterpenes, and carotenoid-derived apocarotenoids originating from β-carotene and ζ-carotene cleavage. (b) Representative phylogeny of Caladeniinae showing major lineages and pollination strategies. Phylogenetic relationships, clade sizes and evolutionary changes in terpenoid-rich floral scent are detailed in Figure 6b. (c) Conceptual model illustrating potential genetic mechanisms underlying the floral volatile transi ions underpinning pollination strategy differences summarised in (b). Relative expression patterns suggest attenuation of selected TPS pathways in sexually deceptive lineages and partial recovery following reversals to food deception, particularly through renewed *TPSb3* expression. Representative food-deceptive species illustrate distinct biochemical outcomes. In *Glossodia major*, coordinated TPS and CCD activities contribute to a complex floral bouquet comprising monoterpenes, sesquiterpenes and volatile apocarotenoids, whereas TPS-b3 and TPS-b4 contribute to monoterpene- and sesquiterpene-rich floral volatiles in *Caladenia denticulata*. The genetic basis of volatile-apocarotenoid variation outside *G. major* remains unresolved. Collectively, the model proposes that changes in TPS expression, catalytic activity and subcellular localisation contribute to floral scent diversification across Caladeniinae, while the evolutionary dynamics of CCD-mediated pathways remain to be established.

## Discussion

### New insights into terpene synthases underpinning floral terpenoid volatile biosynthesis in *Caladenia* orchids

Floral terpenoid diversity in plants is driven by the catalytic versatility and gene family expansions of terpene synthases ^33^, alongside spatiotemporally regulated gene expression ^34,38^. Genomic studies across orchids highlight substantial, lineage-specific TPS expansions that vary across major subfamilies ^15,19,21,39,40^, with recent pan-genomic analyses revealing extensive intrageneric variation ^41^. However, the functional significance of this genomic diversity remains largely unresolved due to a lack of experimental characterisation.

Within the Caladeniinae, *Glossodia major* represents an earlier-diverging lineage (stem age ca. 13 ± 3 Ma) that predates two younger, species-rich radiations (ca. 8 ± 2 Ma), namely subgenus *Calonema* (∼200 spp.) and *Phlebochilus* (∼70 spp.), that employ both food- and sexually deceptive pollination strategies ^42,43^. Characterisation of *GmTPS-b3* (this study), alongside previous functional work on homologs in *Caladenia denticulata Phlebochilus*) and *C. plicata* (subg. *Calonema*) ^20^, reveals that these enzymes act consistently as *α*-pinene synthases lacking detectable sesquiterpene activity. This demonstrates that TPS-b3 monoterpene synthase function has been broadly conserved across Caladeniinae (**Fig. 3, 4, and 5**). Indeed, a similar pattern is observed in TPS7 orthologs of *Aquilegia*, which retain strict specificity for GPP/NPP and consistently produce (+)-limonene while lacking activity toward FPP substrates, despite extensive evolutionary radiation ^44^. Together with consistent dual plastidial and cytosolic localisation in *G. major*, *C. denticulata*, and *C. plicata*, these patterns suggest strong evolutionary constraints maintain specialised monoterpene synthase function across distinct pollination strategies ^26^. This dual localisation suggests that monoterpene biosynthesis may potentially draw on both plastidial and cytosolic precursor pools, consistent with potential inter compartmental exchange of prenyl diphosphates ^45,46^.

In contrast to TPS-b3, TPS-b4 homologs display a distinct evolutionary pattern characterised by catalytic flexibility in some lineages and divergence in others. GmTPS-b4 and CdTPS-b4 retain broad catalytic capacity, utilising multiple substrates to generate diverse mono- and sesquiterpene products **(Fig. 3, 4, and 5)**. Notably, their dominant monoterpene products are consistent with “cineole cassette” TPS enzymes ^47–49^, suggesting retention of key features of this functional class. However, unlike canonical “cineole cassette” TPSs, which typically produce only monoterpenes, both enzymes show substantial catalytic expansion towards sesquiterpene formation. Emerging studies highlight that such multifunctionality is a widespread feature of TPS-b enzymes across plant lineages, e.g. FcoTPS13 and FreTPS15 in *Freesia* ^50^, AjaTPS8 and AoxTPS7in *Aquilegia* ^44,51^, RoTPS059 and RoTPS072 in *Rhododendron* ^52^, CsaTPS5 and CsaTPS32 in *Cannabis* ^53^, and CtrTPS8 in *Cymbidium* ^21^.

In contrast, the homolog CpTPS b4 (CpGES1) in the sexually deceptive *C. plicata* exhibits a narrowed substrate breadth, loss of sesquiterpene forming activity, and plastid specific localisation ^18,20^. This enzymatic specialization supports a highly simplified floral volatile profile dominated by geraniol-derived (S)-citronellol and acetophenones – the active semiochemicals mediating sexual deception. This contrast indicates that lineage-specific constraints can attenuate catalytic promiscuity and subcellular targeting in TPS-b4, driving specialized volatile production within *Caladenia*. Functional divergence among closely related TPS homologs is well documented. In *Freesia*, TPS2 enzymes from several species (e.g. *F. leichtlinii* and *F. corymbosa*) catalyse α terpineol formation from GPP, whereas homologs in closely related species (e.g. *F. viridis* and *F. refracta*) fail to produce this product despite being expressed in the flower ^50^. Divergence can also occur via altered subcellular localization. In *Capsella*, plastid-localized TPS02 drives *β*-ocimene emission in outcrossing *C. grandiflora*, whereas reduced chloroplast targeting diminishes emission in selfing *C. rubella* ^54^.

A complementary pattern of divergence is observed for TPS a1 across the Caladeniinae subtribe. While *GmTPS a1* is strongly expressed and encodes an active sesquiterpene synthase (**Fig. 3, 4, and 5**), its homologs in *C. denticulata* and *C. plicata* are only weakly expressed in highly glandular petal tissue ^18,20^. This progressive reduction in expression from *G. major* to derived *Caladenia* species suggests that regulatory divergence has contributed to the attenuation of TPS a1 function, consistent with the absence of β caryophyllene and α humulene in the respective floral headspace/extracts.

Across multiple plant genera, closely related TPS homologs frequently diverge in catalytic specificity, expression, or subcellular localization despite high sequence identity. Such diversity has been documented in systems including *Costus* (Costaceae) ^55^, *Curcuma* (Zingiberaceae) ^56^, *Mimulus* (Phrymaceae) ^57^, *Freesia* (Iridaceae) ^50,58^, *Aquilegia* (Ranunculaceae) ^44,51^, and *Capsella* (Brassicaceae) ^54^, illustrating the widespread evolutionary flexibility of *TPS* genes underlying floral volatile emission. This evolutionary flexibility is also evident across Caladeniinae, where the broader expression survey supports pollination strategy-associated regulatory divergence in selected TPS genes (**Fig. 6**). Lower *TPSa1* and *TPSb3* expression in sexually deceptive lineages is consistent with attenuation of terpenoid-rich floral scent during their radiation, whereas higher *TPSb3* expression in lineages subsequently shifting to food deception suggests pathway-selective redeployment rather than wholesale restoration of an earlier food-deceptive state. The absence of a comparable pattern for *TPSb4* further indicates that individual TPS modules have followed distinct evolutionary trajectories. Although broader phylogenetically structured sampling of the species-rich *Calonema* and *Phlebochilus* radiations and additional TPS family members is required to test the generality of these patterns, the contrasting trajectories of TPS-b3, TPS-b4 and TPS-a1 reveal how functional conservation, catalytic and compartmental specialisation, and regulatory divergence collectively shape floral terpene evolution across Caladeniinae.

### New insights into carotenoid cleavage dioxygenase underpinning floral apocarotenoids volatile biosynthesis in *Caladenia* orchids

Apocarotenoid diversity in plants arises from the oxidative cleavage of carotenoids by carotenoid cleavage dioxygenases (CCDs), generating a wide range of signalling molecules, pigments, and volatile compounds that contribute to plant development, defence, floral colour, and scent ^35,36^. Despite largely conserved roles among major CCD clades (e.g., CCD1, CCD4, CCD7/8, NCED), apocarotenoid composition varies widely across species, reflecting differences in precursor carotenoid pools and pathway regulation ^59^. In orchids, however, functional characterisation of orchid CCDs has been restricted to a few epiphytic taxa in the subfamily Epidendroideae, including *Oncidium*, *Dendrobium*, *Cymbidium*, and *Phaius* ^60–64^.

In this study, functional characterisation of *Glossodia major* CCDs *in vitro* highlights enzyme specific contributions to floral apocarotenoid production **(Figs. 3, 4, and 5)**. Among these, GmCCD1 uniquely catalyses cleavage of ζ carotene–derived substrates to produce geranylacetone, a major C13 volatile detected in floral headspace. This activity extends to additional substrates and is consistent with canonical C9–C10 (C9′–C10′) cleavage, yielding the corresponding C13 apocarotenoids geranylacetone, β ionone, and pseudoionone from ζ carotene, β carotene, and lycopene, respectively. Additional minor volatile products (e.g. sulcatone) detected in the floral headspace may arise from alternative cleavage events.

Despite biochemical characterisation of CCD1 mediated carotenoid cleavage across several plant genera ^e.g. reviewed^ ^in^ ^59^, direct evidence linking CCD1 activity to floral volatile production remains limited, with only a few characterised examples in species such as petunia, rose, osmanthus, honeysuckle, and clivia ^65–69^. In orchids such as *Dendrobium officinale*, DoCCD1 produces β ionone and pseudoionone from β carotene and lycopene, respectively, but is broadly expressed and not directly linked to floral headspace emission ^62^. The absence of detectable cleavage activity toward phytoene by GmCCD1 in engineered *E. coli* strains accumulating this substrate likely reflects a requirement for extended conjugation in the carotenoid backbone for efficient CCD1-mediated cleavage **(Figs. 3, 4, and 5)**. However, a small subset of highly promiscuous CCD1 enzymes has been reported to accept phytoene in species such as *Rosa damascena*, *Cucumis melo*, and tomato, suggesting a rare but emerging expansion of CCD1 substrate scope^66,70,71^.

Extending beyond CCD1, certain CCD4 variants (e.g. *Vitis vinifera* CCD4b and *Solanum habrochaites* CCD4b) cleave acyclic carotenoids such as ζ-carotene, neurosporene, and lycopene at C9–C10 (or C9′– C10′) positions to yield C_13_ volatiles, including geranylacetone ^72,73^. In contrast, CCD7 enzymes are primarily associated with strigolactone biosynthesis, catalysing 9′–10′ cleavage of 9-cis-β-carotene to generate the C_27_ intermediate 9-cis-β-apo-10′-carotenal ^74^. Although some CCD7s (e.g. AtCCD7) can cleave acyclic carotenoids such as ζ carotene and lycopene, this activity is weak, produces predominantly long chain C_27_ intermediates, and is of unclear biological relevance ^74,75^. Accordingly, *G. major* CCD4 and CCD7 homologs were considered potential contributors to geranylacetone formation; however, no geranylacetone production was detected in engineered *E. coli* strains accumulating phytoene, ζ-carotene, or lycopene **(Figs. 3, 4, and 5)**. Instead, GmCCD4 and GmCCD7a catalysed β-carotene cleavage to produce β-ionone. These findings are consistent with the established role of CCD4 enzymes in generating β-ionone in flowers ^63,69,76–78^ and fruits ^72,79,80^, where β-carotene cleavage contributes to both carotenoid turnover and volatile aroma production.

The detection of pseudoionone in GmCCD7 assays was unexpected, as this product is typically associated with CCD1 activity **(Figs. 3, 4, and 5)**. Although some CCD7 enzymes (e.g. AtCCD7) can cleave acyclic carotenoids such as ζ-carotene and lycopene at the C9–C10 (or C9′–C10′) double bonds, this activity is weak and yields predominantly C_27_ intermediates ^74,75,81^. The observed products therefore likely arise from secondary fragmentation of unstable cleavage intermediates, although an expanded substrate or product scope of GmCCD7 under heterologous expression cannot be excluded. These products were present at low abundance (<1%) relative to geranylacetone, indicating that CCD4 and CCD7 contribute only marginally to the overall floral scent profile.

### Putative roles of *G. major* floral volatiles and multimodal signalling in pollinator attraction

Resolving olfactory signaling in generalized food-deceptive (FD) orchids is challenging due to complex floral bouquets and diverse pollinator guilds, where attraction typically relies on multi-component blends rather than single dominant compounds ^11^. This appears to be the case for *Glossodia major*, whose headspace comprises a dominant carotenoid-derived apocarotenoid (geranylacetone), diverse monoterpenes, and minor sesquiterpenes (notably *β*-caryophyllene; **Fig. 1**). Comparable terpenoid-rich blends occur in related FD orchids, including *Caladenia denticulata* ^20^ and *C. longicauda* ^82^, which, despite representing more derived FD lineages, attract diverse assemblages of generalist pollinators, including native bees, beetles, and wasps ^26,83^.

These key volatiles are widespread floral constituents ^6,84^ that elicit electrophysiological or behavioral responses in generalist insect pollinators ^11^. For instance, *α*-pinene triggers electroantennographic (EAD) activity in *Bombus terrestris* ^85^ and drives behavioural attraction in euglossine bees ^86,87^. Similarly, 1,8-cineole is an EAD-active component attracting euglossine bees in rewarding and deceptive orchids ^85,88^, while *α*-terpineol forms part of EAD-active mixtures attracting bumblebees in other plant families ^89^. Apocarotenoids such as *β*-ionone and geranylacetone trigger strong antennal responses and mediate bee attraction across angiosperms broadly ^5,90,91^. In orchids, the dominant apocarotenoid geranylacetone is frequently reported in floral blends, however, its role in bee attraction remains unconfirmed ^11^. Conversely, minor components like *β*-caryophyllene likely fulfill context-dependent roles within the overall blend ^92–94^.

Crucially, floral signalling across angiosperms operates multimodally, with visual and olfactory cues acting synergistically to mediate pollinator attraction ^95–97^. This integration is well established in deceptive systems globally ^98–104^ and within food- and sexually deceptive Australian orchids specifically ^105–108^. In *G. major*, visual display and volatile emission show striking multimodal alignment: the floral display occupies the bee-blue to bee-blue–green sectors of bee space, matching innate bee preferences and co-flowering rewarding species ^30,109^, while simultaneously emitting a putative bee pollinator-attractive terpenoid blend ^5,11^. A parallel multimodal alignment occurs in the Australian *Thelymitra variegata* complex, where *T. speciosa* matches co-flowering tinsel lilies (*Calectasia* spp.) in both blue–purple floral coloration and monoterpene-dominated scent ^106^.

Rather than reflecting strict mimicry of a specific model, these ubiquitous visual and chemical cues ^6,84^, reinforce generalist bee foraging, consistent with generalized food deception in a shared signalling environment. However, while fruiting success in *G. major* correlates weakly with co-flowering plant density in some seasons ^110^, the precise contribution of floral scent remains unverified. As intense floral colour also enhances background detectability in visually dense communities ^111^, targeted bioassays using synthetic blends and manipulated models of flowers are required to resolve how pollinators integrate these multimodal signals.

## Conclusion

Floral volatile blends are among the most evolutionarily labile traits in flowering plants, yet the molecular mechanisms linking biochemical innovation to floral diversification remain incompletely understood. By integrating floral chemistry, comparative transcriptomics, subcellular localisation, enzyme biochemistry, and evolutionary analyses, we demonstrate that coordinated activity of terpene synthases (TPSs) and carotenoid cleavage dioxygenases (CCDs) underpins the production of monoterpenes, sesquiterpenes, and volatile apocarotenoids in *Glossodia major*. Together with emerging phylogenomic insights, our findings suggest that major transitions in floral phenotype, pollination strategy, and scent chemistry have accompanied the diversification of the subtribe Caladeniinae (**Fig. 7**). Functional comparisons among *G. major* and closely related *Caladenia* species (**Fig. 7a**) reveal a mosaic of evolutionary conservation, divergence, and innovation across TPS mediated pathways. We therefore propose a conceptual model (**Fig. 7b**) in which repeated modification of these pathways, through changes in catalytic function, gene expression, and subcellular partitioning, contributed to the reduction, specialisation and re-emergence of floral volatile traits across lineages (**Fig. 7c**). More broadly, our results support a model of evolutionary tinkering in which conserved biosynthetic genes are repeatedly modified and differentially deployed to generate phenotypic diversity^113^, providing candidate mechanistic links between molecular evolution, pollinator-mediated selection, and major shifts in floral scent across flowering plants.

## Materials and Methods

### Orchid sampling

Flowering individuals of *Glossodia major* were sampled at Black Mountain Nature Reserve (Canberra, Australia) during peak flowering. Floral material was collected from more than 100 randomly selected purple flowered individuals under clear, warm midday conditions, when flowers were fully open and floral scent emission was most intense (to the human nose). Flowers with intact stems were excised at the base in the field, placed immediately into water filled insulated containers for transport, and transferred to the laboratory for immediate analyses.

### Floral headspace sampling and analysis

Floral volatile headspace was sampled from four bouquets (24, 25, 27, and 36 flowers) enclosed in heat-resistant oven bags, with cut stems in water-filled 50 mL tubes. Two parallel controls of cut non-floral stems were prepared to account for background emissions. After 30 min equilibration, air was drawn at ∼200 mL min⁻¹ through pre-cleaned Porapak Q adsorbent tubes positioned ∼4 cm from flowers. Collections were conducted over three consecutive 24-h intervals. Trapped volatiles were eluted with 1.5 mL dichloromethane containing tridecane (0.5 ng µL⁻¹) as an internal standard and stored at −20 °C.

Samples were analyzed on an Agilent 8860 GC coupled to an Agilent 5977B MSD with a DB-FFAP capillary column (30 m × 0.25 mm × 0.5 µm). Aliquots (1 µL) were injected splitless at 250 °C with He carrier gas (1.2 mL min⁻¹). The oven program held at 40 °C (1 min), ramped at 10 °C min⁻¹ to 230°C, and held for 10 min. Mass spectra were acquired at 70 eV (m/z 40–450). Retention indices (RI) were determined using n-alkanes (C_8_ – C_44_; 10 ng µL⁻¹). 1,8-cineole and α-terpineol (Sigma-Aldrich) were run as authentic standards. Data were processed in Agilent MassHunter (v10.0), matching spectra and RI against NIST17 (https://www.nist.gov/) using the webchem R package ^112^. Compounds with <80% spectral match or >10 RI difference from literature were annotated as unknown. Relative abundances were calculated from integrated peak areas after subtracting control signals (Please see Supplementary Methods for further detail).

### Transcriptome sequencing & analysis

Publicly available RNA-seq datasets spanning petal, stem, and leaf tissues of common purple-flowered *G. major* ^30^ were re-analyzed to examine reproductive tissue-specific gene expression. Differential expression (petal vs. leaf/stem) was evaluated using edgeR with a quasi-likelihood F-test ^114^. Genes with FDR < 0.01 were considered significantly differentially expressed, reported as FPKM. Expression overlaps were evaluated using UpSet plots in TBtools ^115^. Functional enrichment of MapMan4 BIN categories ^116^ was assessed via a hypergeometric test with FDR correction < 0.05 (Please see Supplementary Methods for further detail).

### Terpene synthase and carotenoid cleavage dioxygenase gene family analysis

Translated protein sequences from the G. major transcriptome were screened for TPS and CCD family members using HMMER v3 ^117^. Hidden Markov Models corresponding to conserved TPS N-terminal (PF01397) and C-terminal (PF03936) domains and the CCD carotenoid oxygenase domain (PF03055) were used for candidate identification. Candidates were aligned with functionally characterized plant proteins using MAFFT v7 ^118^ and manually curated across catalytic motifs. Maximum-likelihood phylogenies were constructed in IQ-TREE v2 ^119^ using ModelFinder ^120^ and 1,000 ultrafast bootstrap replicates (Hoang et al., 2018), then visualized in iTOL ^121^ with functional annotations mapped (Please see Supplementary Methods for further detail).

### Isolation and construction of terpene synthase and carotenoid cleavage dioxygenase expression plasmids

Codon-optimized full-length CCD and TPS coding sequences were synthesized (CWBio, China) and amplified using gene-specific primers (**Supplementary data S10**) with Phanta Max Master Mix (Vazyme, China). Full-length constructs were cloned into pCNHP-YFP. TargetP 2.0 (https://services.healthtech.dtu.dk/services/TargetP-2.0/) was used to predict chloroplast transit peptides, and corresponding N-terminally truncated sequences were cloned into pMAL-c5X. Constructs were assembled via homologous recombination using ClonExpress Ultra V3 (Vazyme, China) for subsequent subcellular localization and enzymatic activity assays (Please see Supplementary Methods for further detail).

### Agrobacterium-mediated transient expression in *Nicotiana benthamiana*

*Nicotiana benthamiana* plants were grown under controlled conditions (26 °C, 60% RH, 14 h light/10 h dark, 100 µmol m⁻² s⁻¹). Four-week-old plants were used for *Agrobacterium tumefaciens* (strain GV3101 carrying pCNHP-YFP) transient expression ^122^. Exponential phase cultures were harvested, resuspended in infiltration buffer (10 mM MgCl₂, 10 mM MES, pH 5.6, 100 µM acetosyringone) to OD_600_=0.8, and incubated in the dark for 2 h. Suspensions were infiltrated into abaxial leaf surfaces, and plants were maintained for 3 days prior to imaging (Please see Supplementary Methods for further detail).

### Subcellular localization

Full-length coding sequences were cloned into pCNHP-YFP and introduced into *A. tumefaciens* GV3101 for transient expression in *N. benthamiana* leaves as described above. Fluorescence signals were observed directly from the abaxial epidermal cells using a Zeiss LSM 880 inverted confocal laser scanning microscope. YFP fluorescence was detected at 520-550 nm following excitation at 514 nm, and chlorophyll autofluorescence at 680-720 nm following excitation at 633 nm.

### Carotenoid cleavage dioxygenase (CCD) enzyme assays

Functional characterization of CCDs was performed using *E. coli* BL21(DE3) strains engineered to accumulate phytoene (pAC-PHYT), ζ-carotene (pAC-ZETA), lycopene (pAC-LYC), or β-carotene (pAC-BETA) ^37^. Recombinant strains were grown in LB medium (30 C) to OD_600_≈0.6, induced with 0.01 mM IPTG, and incubated at 28 °C for 12 h. Cells were harvested, extracted with MTBE, concentrated via vacuum centrifugation, and analyzed by GC–MS (Please see Supplementary Methods for further detail).

### *In vitro* terpene synthase enzyme assays

N-terminally truncated TPS sequences (lacking predicted transit peptides) were cloned into pMAL-c5X and transformed into *E. coli* Rosetta2 (DE3) pLysS (Novagen, USA). Expression was induced with 0.1 mM IPTG at 16 °C for 16 h (OD_600_≈0.4). Cells were lysed by sonication in buffer (20 mM Tris–HCl pH 7.4, 200 mM NaCl, 1 mM EDTA, 1 mM DTT, 1 mM PMSF). MBP-fusion proteins were purified using amylose resin (Yeasen, China) and verified by SDS–PAGE. Assays were conducted at 30 °C for 1 h using 50 µM prenyl diphosphate substrates (GPP, NPP, E,E-FPP, Z,Z-FPP, GGPP; Echelon Biosciences) following established methods ^20,122^. Reactions were extracted with MTBE, concentrated under N_2_, and analyzed by GC–MS. MBP-tag only reactions served as negative controls (Please see Supplementary Methods for further detail).

### Gas chromatography-mass spectrometry analysis of enzyme assay products

GC–MS analysis was performed essentially as described previously ^20^. Enzymatic products were analyzed on a Shimadzu QP-2020NX GC–MS system fitted with an Rxi-5Sil column (30 m × 0.25 mm × 0.25 µm; Restek). Aliquots (1 µL) were injected splitless at 250 °C with He carrier gas (1 mL min⁻¹). The oven program was: 40 °C (3 min), ramped at 10 °C min⁻¹ to 280 °C (5 min), then at 20 °C min⁻¹ to 300 °C (2 min). Compounds were identified by comparing retention times and mass spectra against authentic standards (MedChemExpress) and the NIST20 library (Please see Supplementary Methods for further detail).

## Author contributions

DCJW and FZ designed the study. YNZ, JP, WZ, DCJW performed the research and analyzed the data. DCJW, EP, FZ, and RP secured funding. DCJW and FZ wrote the article with assistance from all authors.

## Acknowledgements

The pAC-ZETA and pAC-PHYT vector were kindly shared by Professor Li-En Yang (Jiangsu Marine Fisheries Research Institute, Jiangsu, China). We thank the Australian Capital Territory Environment, Planning and Sustainable Development Directorate for permission and permits (Permit no.: PL201912) to obtain plant material from Black Mountain Nature Reserve (Canberra, Australia). This work was supported by the Australian Research Council projects DE190100249 to DW and DP210100471 to RP and EP, the National Natural Science Foundation of China (32300230 to FZ), the Natural Science Foundation of Jiangsu Province (BK20230785 to FZ), and an AGRTP scholarship to JP. D.C.J.W. also acknowledges financial support from an Adelaide University Future Making Fellowship.

## Data availability

The raw RNA-seq data presented in this study can be found in NCBI Sequence Read Archive (http://www.ncbi.nlm.nih.gov/sra) under the SRA study and BioProject accessions SRP471419 and PRJNA1039539, respectively. Full-length coding sequences of *Glossodia major* TPS-b3 (PZ775651), TPS-b4 (PZ775650), TPS-a1 (PZ775649), CCD1 (PZ775645), CCD4 (PZ775646), CCD7a (PZ775647), and CCD7b (PZ775648) have been deposited in NCBI GenBank (https://www.ncbi.nlm.nih.gov/genbank/).

